# Computational hemodynamic indices to identify Transcatheter Aortic Valve Implantation degeneration

**DOI:** 10.1101/2024.02.09.579647

**Authors:** Luca Crugnola, Christian Vergara, Laura Fusini, Ivan Fumagalli, Giulia Luraghi, Alberto Redaelli, Gianluca Pontone

## Abstract

**Purpose:** Structural Valve Deterioration (SVD) is the main limiting factor to the long-term durability of bioprosthetic valves, which are used for Transcatheter Aortic Valve Implantation (TAVI). The aim of this study is to perform a patient-specific computational analysis of post-TAVI blood dynamics to identify hemodynamic indices that correlate with a premature onset of SVD.

**Methods:** The study population comprises two subgroups: patients with and without SVD at long-term follow-up exams. Starting from pre-operative CT images, we created reliable post-TAVI scenarios by virtually inserting the bioprosthetic valve (stent and leaflets), and we performed numerical simulations imposing realistic inlet conditions based on patient-specific data. The numerical results were post-processed to build suitable synthetic scores based on normalized hemodynamic indices.

**Results:** We defined three synthetic scores, based on hemodynamic indices evaluated in different contexts: on the leaflets, in the ascending aorta, and in the whole domain. Our proposed synthetic scores are able to clearly isolate the SVD group. Notably, we found that leaflets’ OSI individually shows statistically significant differences between the two subgroups of patients.

**Conclusion:** The results of this computational study suggest that blood dynamics may play an important role in creating the conditions that lead to SVD. Moreover, the proposed synthetic scores could provide further indications for clinicians in assessing and predicting TAVI valves’ long-term performance.

## 1 Introduction

Transcatheter Aortic Valve Implantation (TAVI) is a minimally invasive technique for the treatment of severe Aortic Stenosis (AS). It consists in the deployment of a stented bioprosthetic valve inside the native aortic valve via a percutaneous catheter, restoring a physiological function of the aortic orifice. Introduced in 2002 as an alternative to open-heart Surgical Aortic Valve Replacement, TAVI has become the standard of care for patients with severe AS at prohibitive surgical risk, and the preferred treatment for many intermediate and high-risk elderly patients [1, 2]. Moreover, the results of the recent PARTNER 3 and Evolut Risk clinical trials suggest that TAVI might be the preferred option for AS treatment even in low-risk younger patients [3–6].

Assessing the long-term durability of TAVI valves is of utmost importance in order to extend the TAVI procedure to younger patients with longer life expectancy. However, TAVI is a fairly new technique, thus there is a lack of long-term follow-up data; moreover, available data are mostly related to older generation devices implanted on elderly patients, having limited life expectancy [7]. Therefore, the prediction of TAVI effectiveness and the understanding of the mechanisms underlying degeneration over the years are nowadays still challenging issues.

Structural Valve Deterioration (SVD) leading to regurgitation or stenosis is the main limiting factor to the long-term durability of TAVI bioprosthetic valves [8, 9]. This is an irreversible process manifested by gradual degenerative changes in the bioprosthesis, such as pannus growth, leaflet fibrosis and calcification, possibly leading to ruptures and perforations of the leaflets [10]. Recent studies provided evidence that multiple processes are involved in SVD pathogenesis, including immune rejection and atherosclerosis-like tissue remodeling [9, 10], suggesting a possible influence of aortic hemodynamics on the development of SVD [11].

Computational models have been extensively employed within the TAVI framework. A structural analysis, via the Finite Element Method, has been usually applied to study the implantation of the bioprosthetic valve, assessing the impact of device design and anatomical features on the outcome of the intervention [12–14]. Post-TAVI hemodynamics have been numerically simulated using Computational Fluid Dynamics (CFD) or Fluid-Structure Interaction approaches, usually investigating complications such as para-valvular leakage, embolism risk, and thrombogenicity [15–17]. The investigation in [11] represents the first computational hemodynamic study focusing on SVD, where a few patient-specific cases were analysed to explore possible relations between hemodynamics shortly after the TAVI procedure and the long-term degeneration of the bioprosthetic valves. In particular, the authors correlated the early post-TAVI stress distribution on the proximal aortic wall with the presence of SVD detected at 5-10 years follow-up exams.

In the current work we delve deeper into this analysis, overcoming the main limitations of [11], which are: i) the focus on the sole systolic phase; ii) the reduced number of patients analyzed; iii) the absence in the geometric model of the bioprosthetic valve’s leaflets; iv) the use of the same boundary conditions for each patient. This allows us to propose reliable hemodynamic indices in early post-TAVI stages that correlate with the premature onset of SVD. In particular, the main objectives and novelties of the work can be summarized as follows:

- A complete computational-hemodynamics investigation of fourteen patients, comprising the whole heartbeat (systole and diastole), the presence of the bioprosthetic valve’s leaflets, and a turbulence model;
- The prescription of a realistic inlet flow rate condition tuned on patient-specific post-operative cardiac output measurements;
- The definition of new hemodynamic-based synthetic scores, obtained by combining different indices that are easily computable as post-processing of the numerical simulations, able to discriminate between patients with and without SVD at 5-10 years follow-up exam.

To this aim, we start from pre-operative Computed Tomography (CT) images and we perform a virtual implantation of the bioprosthetic valve to obtain trustworthy early post-TAVI computational scenarios, representative of few days after the intervention.

The analyses of this work are strongly based on the assumption that hemodynamics in early post-TAVI stages can influence the long-term degeneration of the implanted valves. This can be motivated by the creation of specific hemodynamic conditions which trigger a cascade of events possibly contributing to deterioration [10].

## 2 Materials and Methods

In this work we present a retrospective computational study aiming at finding hemodynamic indications about Structural Valve Deterioration (SVD). This encompasses the following steps:

1. Virtual implantation of a bioprosthetic valve inside patient-specific aortic root geometries, reconstructed from pre-operative CT scans;
2. Patient-specific CFD analysis of early post-TAVI hemodynamics, performed in the virtual scenarios obtained after step 1., when the bioprosthetic valve is yet to be degenerated;
3. Post-processing of the CFD results to identify hemodynamic indices that correlate with the presence of SVD detected at 5-10 years follow-up exam.

This is done with the prospective aim of employing our model in a clinical setting, in order to predict TAVI valves’ degeneration and provide indications for a personalized follow-up planning. Analogously to what is done by companies such as FEops (https://www.feops.com/) and HeartFlow (https://www.heartflow.com/), we introduce in our model some simplifications in order to reduce the computational cost of the numerical simulations:

- Cylindrical shape of the virtually implanted stent is assumed and no deployment simulation is performed (see Section 2.2);
- The bioprosthetic leaflets dynamics is described in an on/off modality and no Fluid-Structure Interaction is accounted for (see Section 2.2);
- Rigid aortic walls are assumed (see Section 2.3).
- Suxh simplification will be discussed in Section 4.

In what follows we present the available clinical data (Section 2.1) and the generation of patient-specific early post-operative virtual scenarios (Section 2.2); then, we describe the mathematical and numerical models for aortic hemodynamics in presence of the TAVI bioprosthetic valve (Section 2.3), and we discuss the prescription of inlet boundary conditions (Section 2.4); finally, we introduce the analyzed hemodynamic indices and we build synthetic SVD-discriminating scores (Section 2.5).

### 2.1 Clinical Data

Seven cases of patients who underwent TAVI between 2008 and 2012 with SVD at longterm follow-up were identified and seven cases without SVD at long-term follow-up were randomly extracted from the same cohort and matched for baseline characteristics. We will refer to the former group as DEG patients and to the latter one as NODEG patients. The mean follow-up time is 7.2 *±* 1.9 years. The definition of SVD was SVD stage 3 as previously defined [18]. The study was approved from IRB of Centro Cardiologico Monzino and registered with number R1264/20.

In this study we consider only patients who received an Edwards SAPIEN balloon-expandable valve with size 23, which is the most likely to develop SVD between the Edwards SAPIEN valves [3]. In particular, it consists of a trileaflet valve made of bovine pericardium mounted on cobalt-chromium stent with an external diameter of 23 *mm* and a height of 14.5 *mm*. On the ventricular side of the stent frame, an inner polyethylene terephthalate fabric skirt is applied [11, 19] (see Figure 1a).

**Fig. 1:**
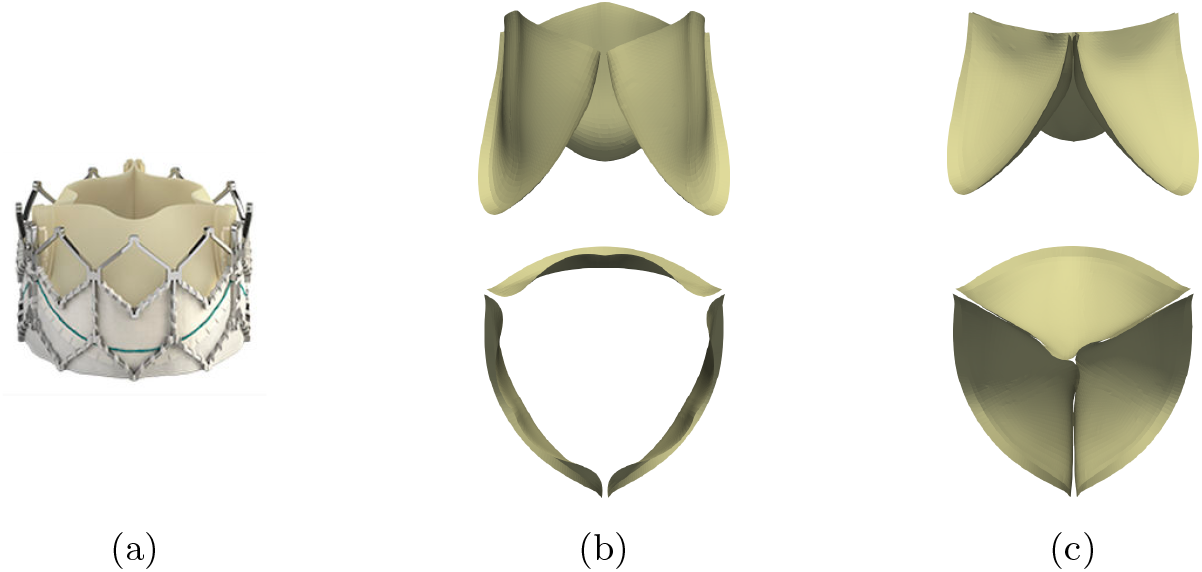
(a) Edwards SAPIEN bioprosthetic valve (source *Medical Expo*: https://www.medicalexpo.com/). (b)-(c) Trileaflet models obtained in [20] from a structural numerical simulation: (b) open configuration, (c) closed configuration

For each patient we have at disposal:

- a pre-operative CT scan taken at late diastole with a Discovery HD750 scanner (GE Healthcare) using the following configuration: 64*×*0.625 *mm*; gantry rotation time 350 *ms*; tube voltage, 120 *kV p*; and effective tube current, 650 *mA*. Contrast enhancement was achieved with a triphasic injection of an 80 *ml* bolus of Iomeron 400 *mg/ml* (Bracco Imaging S.p.A.) through an antecubital vein at a 5 *ml/s* infusion rate, followed by 50 *ml* of saline solution, and a further 50 *ml* bolus of contrast at 3.5 *ml/s* [3].
- early post-TAVI Transthoracic Echocardiography (TTE) data obtained with commercially available equipment (iE33 or Epiq, Philips Medical System, or Vivid-9, GE Healthcare) [3] between 2 days and 7 months after the implantation. Specifically, in this work we exploit patient-specific cardiac output measurements obtained from TTE data (see Section 2.4).

### 2.2 Post-operative virtual domain generation

We build post-TAVI computational domains starting from pre-operative CT scans. The procedure is carried out using the Vascular Imaging Toolkit (VMTK, http://www.vmtk.org [21]) and Paraview (https://www.paraview.org) software, and comprises the following steps:

1. Reconstruction of the aortic root geometry and calcium deposits from the pre-operative CT scan;
2. Virtual implantation of the bioprosthetic valve’s stent inside the pre-operative reconstructed aortic root;
3. Positioning of suitable leaflets geometries, taken from [20], inside the virtually implanted stent;
4. Generation of the volumetric mesh.

The first two steps of the procedure are detailed in [11]. In particular, the virtual stent, modelled as a hollow cylinder, is oriented using the centerline of the reconstructed pre-operative aortic geometry and positioned at the barycenter of the aortic annulus. The virtual stent is then rigidly translated along its radial direction to account for the patient-specific calcification pattern on the native aortic valve and the aortic annulus is possibly deformed in case of a dimensional mismatch with the bioprosthetic valve, analogously to what happens during TAVI due to the balloon inflation. The resulting computational domain is shown for one patient in Figure 2a and comprises: the aortic wall downstream the aortic annulus, the interface between blood and the aortic annulus, the interface between blood and the stent, an inlet section and a outlet section.

**Fig. 2:**
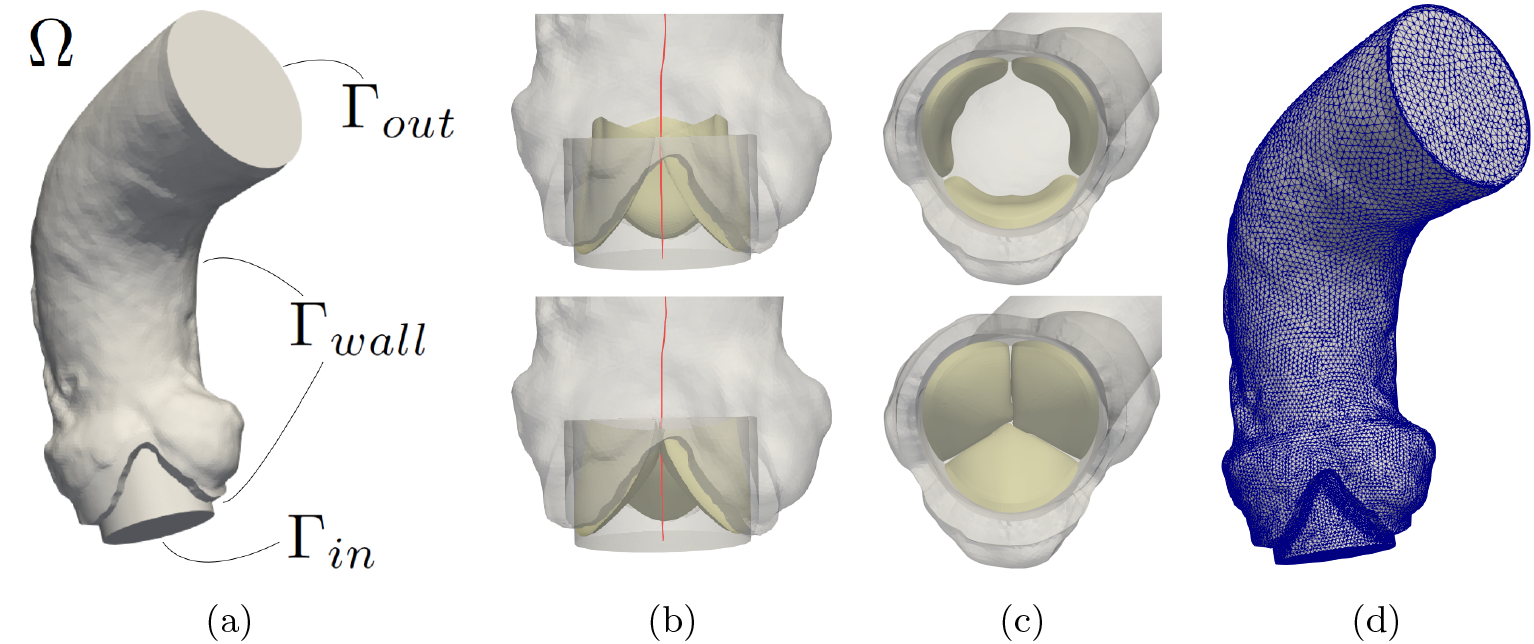
(a) Computational domain Ω for one patient. Γ_*in*_ is the inlet boundary; Γ_*out*_ is the outlet boundary; Γ_*wall*_ is the wall, composed by the aortic wall, the interface between blood and the aortic annulus and the interface between blood and the bioprosthetic valve’s stent. (b) Bioprosthetic leaflets positioned inside the virtually implanted stent using the centerline of computational domain (top: open configuration, bottom: closed configuration). (c) Orientation of the bioprosthetic leaflets in agreement with the orientation of the native leaflets (top: open configuration, bottom: closed configuration). (d) Volumetric computational mesh with a boundary layer close to the aortic wall and a local mesh refinement close to the bioprosthetic leaflets.

One of the major novelties of this work with respect to [11] is the introduction of the bioprosthetic valve’s leaflets (step 3 in the list above). The leaflets geometries, in open and closed configurations (Figure 1b-1c), were obtained in [20] starting from an idealized trileaflet model and applying physiological pressure gradients in a structural numerical simulation. The leaflets are positioned inside the virtually implanted stent by exploiting the centerline of the generated computational domain (see Figure 2b). In particular, we orient the bioprosthetic leaflets in order to match the native ones (see Figure 2c), in accordance with common clinical practice [22]. Note that the thickness of the leaflets will be defined directly inside the mathematical model (see Section 2.3). The open and closed configurations of the leaflets are used to provide a quasi-static on/off modeling of opening/closure dynamics. Specifically, we switch between these two rigid configurations in agreement with the flow rate profile imposed at the inlet (see Section 2.4), which identifies the systolic and diastolic phases.

Finally, a volumetric mesh is generated inside the computational domain (see Figure 2d). For all fourteen patients, tetrahedral meshes were generated using VMTK with an average mesh size of approximately 1 *mm*, a local refinement of 0.25 *mm* close to the bioprosthetic valve’s leaflets and a boundary layer composed by three layers close to the aortic wall (finest layer’s thickness = 0.14 *mm*). The mesh size is chosen after a mesh convergence analysis performed for patient NODEG2 and based on sequential refinements. In particular, we stop the refinement when the mean value in time 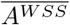 of the area of the aortic wall with Wall Shear Stress (WSS) greater than 5 *Pa* (see Section 2.5) shows a relative difference below 1%, as a consequence of a refinement factor *ℛ* = 0.77 (that is the ratio of two consecutive mesh sizes is equal to 0.77). We identify two characteristic sizes for our meshes, the maximum mesh size *h*_*max*_ and the boundary layer’s finest layer thickness *h*_*BL*_; Table 1 shows the results of the convergence analysis: *h*_*max*_ = 1.75 *mm* and *h*_*BL*_ = 0.14 *mm* are selected using the considered criterion.

**Table 1:** Mesh convergence analysis: *i* is the refinement step; 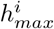 is the maximum mesh size; 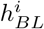 is the boundary layer’s finest layer thickness; 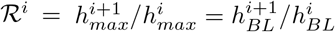 is the refinement factor; 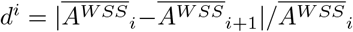 is the absolute relative difference.

| $i$ | $h_{max}^i$ | $h_{BL}^i$ | $\mathcal{R}^i$ | $d^i$ |
| --- | --- | --- | --- | --- |
| 0 | 2.95 mm | 0.23 mm | 0.77 | 12.78% |
| 1 | 2.27 mm | 0.18 mm | 0.77 | 4.35% |
| 2 | 1.75 mm | 0.14 mm | 0.77 | 0.61% |
| 3 | 1.35 mm | 0.11 mm |  |  |

### 2.3 Mathematical and numerical methods

We are interested in studying aortic blood dynamics in presence of the TAVI bioprosthetic valve. We model blood as an incompressible homogeneous Newtonian fluid, as it is common practice in large vessels like the aorta [23], thus describing its dynamics with the Navier-Stokes equations.

Our computational domain is the one resulting from the post-operative virtual domain generation procedure described in Section 2.2 (Figure 2a). In particular, the aortic wall and the interfaces between blood and both the stent and the aortic annulus are treated as rigid boundaries. A flow rate condition is imposed at the inlet boundary, whereas a physiological pressure *P*_*Wig*_(*t*), representing a Wiggers aortic pressure (Figure 3a) profile, is imposed at the outlet boundary.

**Fig. 3:**
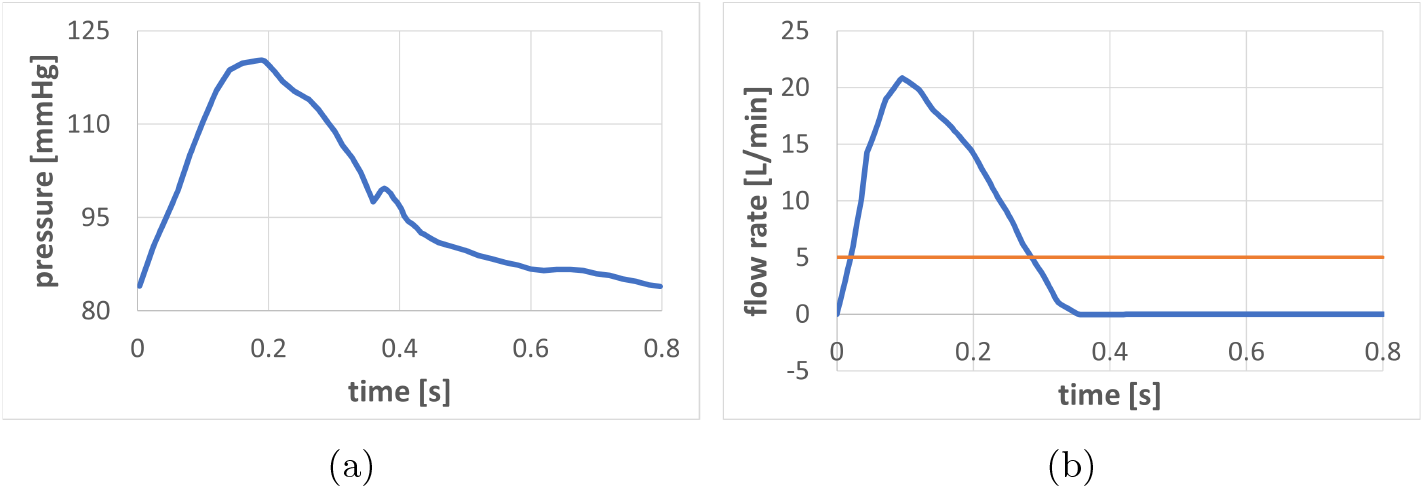
(a) Wiggers aortic pressure profile imposed at outlet section. (b) Physiological flow rate waveform (mean value: 5 *L/min*) used to impose inlet boundary conditions. Patient-specific mean flow rate values are recovered for each patient according to the cardiac output measurements reported in Table 2.

We account for the presence of turbulence inside the ascending aorta [24] by using the Large Eddy Simulation (LES) *σ*-model [25–27], thus adding a viscous term which accounts for the non-resolved scales. The bioprosthetic valve’s leaflets are implicitly represented as an obstruction to the flow using the Resistive Immersed Implicit Surface (RIIS) method [28, 29], thus adding to the momentum equation of the Navier-Stokes system a local penalization term, which represents the adherence of the blood to the leaflets.

The resulting strong formulation of the considered mathematical model reads: Given the initial blood velocity **u**_0_, for each time *t >* 0, find the blood velocity **u** and the blood pressure *p*, such that:

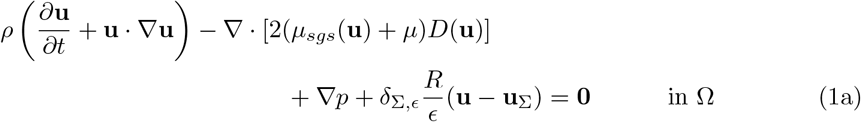

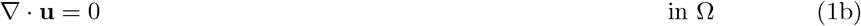

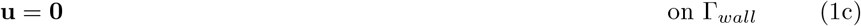

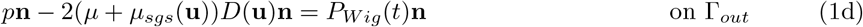

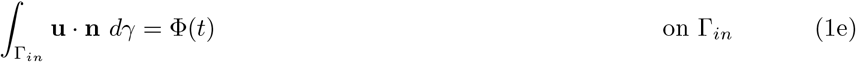

where *ρ* = 1060 *kg/m*^3^ and *µ* = 3.5 *×* 10^−3^ *Pa · s* are the blood density and viscosity, respectively, and *µ*_*sgs*_ is the sub-grid scale viscosity introduced by the LES *σ*-model,which depends on the velocity field [25];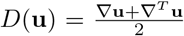, where the apex *T* stands for transpose, is the strain rate tensor; the last term of equation (1a) represents the RIIS method: Σ is the bioprosthetic leaflets surface, *R* = 10^5^ *Kg/m · s* represents a resistance coefficient, *ϵ* = 0.2 *mm* is the leaflets’ half-thickness, *δ*_Σ,*ϵ*_ is a smoothed Dirac delta function representing a layer with thickness 2*ϵ* around Σ (thus we consider 0.4 *mm* thick leaflets) and **u**_Σ_ is the velocity of the bioprosthetic leaflets. According to the on/off modeling used in this work for the valve dynamics (see Section 2.2), we set **u**_Σ_ = **0**. The choice of the prescribed flow rate Φ(*t*) will be discussed in Section 2.4.

Being a defective condition, equation (1e) is not enough to guarantee existence and uniqueness of a solution. In order to prescribe such condition we exploit an augmented weak formulation of the Navier-Stokes problem based on Lagrange multipliers and on the assumption that the traction on Γ_*in*_ is normal to the surface and constant in space, as proposed in [30, 31]. This approach allows us to impose the desired flow rate profile at the inlet boundary without fixing a spatial profile for the velocity field, which would highly influence the results of the numerical simulations.

For the time discretization of the continuous problem we consider a first order implicit backward Euler scheme with an explicit treatment of the convective field and the sub-grid scale viscosity *µ*_*sgs*_(**u**). We consider a time-step Δ*t* = 10^−3^ *s* chosen after performing a sensitivity analysis, analogous to the one for the mesh size (see Section 2.2). For space discretization we employ piece-wise linear finite elements for the velocity and pressure fields with a Streamline Upwind Petrov-Galerkin/Pressure Stabilizing Petrov-Galerkin (SUPG/PSPG) stabilization scheme [27, 32, 33]. At the outlet boundary we add a backflow stabilization, as proposed in [34], to prevent instabilities due to the artificial cut of the computational domain.

The mathematical and numerical methods presented above are implemented in the multi-physics high-performance library life^x^ [35, 36] (https://gitlab.com/lifex, https://doi.org/10.5281/zenodo.7852088), developed at MOX, Dipartimento di Matematica, with the collaboration of LaBS, Dipartimento di Chimica, Materiali ed Ingengneria Chimica (both at Politecnico di Milano).

In order to assess the LES model quality we consider the Pope criterion [37], namely we compute the fraction of turbulent kinetic energy in the resolved scales *M* (**x**, *t*):

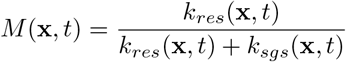

where *k*_*res*_(**x**, *t*) is the resolved turbulent kinetic energy per unit mass and *k*_*sgs*_(**x**, *t*) = *µ*_*sgs*_(**x**, *t*)^2^*/*(*C*Δ*ρ*)^2^ [38] is the sub-grid scale turbulent kinetic energy, with *C* = 1.5 a LES constant [25] and Δ = 1 *mm* a representative average value of the mesh size. Notice that a good LES model should be able to resolve at least 80% of the total turbulent kinetic energy [37, 39]. We analyzed the same patient considered for the mesh convergence and we found that the fraction of volume in the computational domain where *M >* 0.8 is always greater than 88%, with mean value during the heartbeat of 93%. Thus we deem our discretization strategy as sufficiently refined for the considered LES model.

We numerically simulate 6 complete heartbeats and we discard the first one to remove the influence of the null initial condition.

Simulations were run in parallel on 56 cores with CPU’s Xeon E5-2640v4@2.4 GHz, using the computational resources available at MOX, Dipartimento di Matematica, Politecnico di Milano.

### 2.4 On the prescription of inlet boundary conditions

The aim of this work is to exploit the mathematical and numerical methods presented in Section 2.3 to provide a reliable description of blood-dynamics in early post-TAVI scenarios. To this aim, a suitable flow rate Φ(*t*) should be selected in the inlet boundary condition (1e). Specifically, we impose the physiological waveform taken from [40] (see Figure 3b) and, for each patient, we adapt its magnitude to obtain patient-specific mean flow rate values that match early post-TAVI TTE cardiac output measurements (see Table 2). Notice that, in absence of any information regarding the patients’ Heart Rate (HR), we assumed for all of them the same value HR = 75 *bpm*.

**Table 2:** Cardiac output measurements obtained from early post-TAVI TTE data and used to adapt the waveform in Figure 3b in order to impose for each patient the patient-specific mean flow rate value at the inlet section.

| ID | CO [L/min] | ID | CO [L/min] |
| --- | --- | --- | --- |
| **DEG1** | 5.90 | **NODEG1** | 5.35 |
| **DEG2** | 6.05 | **NODEG2** | 4.50 |
| **DEG3** | 5.45 | **NODEG3** | 5.45 |
| **DEG4** | 4.85 | **NODEG4** | 5.10 |
| **DEG5** | 5.45 | **NODEG5** | 4.35 |
| **DEG6** | 5.95 | **NODEG6** | 5.85 |
| **DEG7** | 5.30 | **NODEG7** | 5.05 |

#### Hemodynamic indices

We aim to discriminate between DEG and NODEG patients, thus we perform a post-processing of the numerical results obtained with the model presented in Section 2.3-2.4 Specifically, we analyze the following hemodynamic indices:

##### Wall Shear Stress

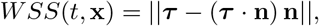

where **n** represents the unit normal vector to a given surface and *τ* =2 (*µ*_*sgs*_(**u**) + *µ*) *D*(**u**)**n** is the traction. Additionally, we consider *A*^*W SS*^(*t*), the area of the aortic wall where *WSS* is greater than the threshold of 5 *Pa*. This threshold identifies regions of high WSS, not necessarily pathological. Notice that instantaneous WSS magnitude typically ranges from 1 to 7 [*Pa*] in straight vessels of the arterial system [41]. High values of WSS can be associated with changes in the endothelial cell behavior, potentially exacerbating inflammation [42];

##### Q-criterion

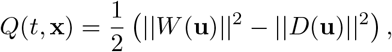

where 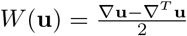 is the vorticity tensor. Positive values of this index identify regions where vorticity dominates over laminar friction. Additionally, we consider 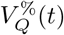, the fraction of volume where *Q* is greater than a selected threshold [43] of 50000 *Hz*^2^. This threshold is chosen to properly isolate vortical structures. The proposed volume fraction gives a measure of flow disturbance inside the considered domain. Disturbed flow has a well-proven impact on vascular endothelial cells and contributes to the pathophysiology of clinical conditions such as in-stent re-stenosis as well as aortic valve calcification [44];

##### Diastolic Time-Averaged Wall Shear Stress

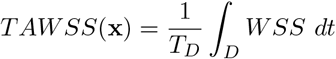

where *D* is the diastolic interval and *T*_*D*_ its duration. Additionally, we consider *avgTAWSS*, the average-in-space *TAWSS* on the aortic face of the bioprosthetic leaflets. Notice that low shear stresses on the aortic side of the aortic valve can enhance the permeability of the endothelial cell barrier with respect to small molecules [45];

##### Oscillatory Shear Index

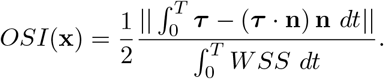

Additionally, we consider *avgOSI*, the average-in-space *OSI* on the aortic face of the bioprosthetic leaflets. Oscillatory shear stresses on the aortic side of the valve promote inflammation and calcification [46].

In order to discriminate between DEG and NODEG patients, we assign a score related to each hemodynamic index and we combine them to define SVD-discriminating synthetic scores. In particular, indicating with the bar average values over time, we consider the scalar quantities 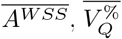, *avgTAWSS, avgOSI* and we normalize them exploiting the inter-patient variability. We split the study population (14 patients) into a normalization set (12 patients) and a blind set (2 patients: DEG7 and NODEG7), we compute the first (*ψ*_1_) and third (*ψ*_3_) quartiles of each score in the normalization set and we linearly project the interval [*ψ*_1_ − 1.5 ∗*IQR, ψ*_3_ + 1.5 ∗*IQR*] into the interval [−1, 1], where *IQR* = *ψ*_3_ − *ψ*_1_ is the inter-quartile range. So that, four normalized scores, related to the proposed hemodynamic indices, are assigned to each patient.

The SVD-discriminating scores are defined as a linear combination of the mean and maximum values of these four scores. In particular, by splitting the hemodynamic indices related to the aorta and the ones related to the leaflets, we consider:

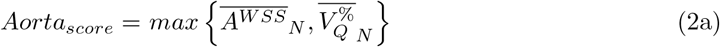

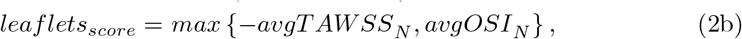

where the subscript *N* identifies the normalized scores and the opposite of *avgTAWSSN* is considered in order to associate low values of this hemodynamic score to high values of the synthetic score. Finally, we introduce a global SVD-discriminating score that synthesizes each considered index:

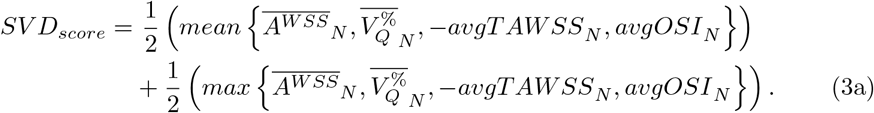

These scores include, in a synthetic way, multiple hemodynamic features that may influence the degeneration of TAVI valves. This is an accordance with our belief that analyzing such features individually could not account for the complexity of the SVD process.

## 3 Results

### 3.1 On the accuracy of the virtual stent implantation

The early post-TAVI computational domains obtained following the procedure presented in Section 2.2 are reported in Figure 4 together with the positioned bioprosthetic leaflets in open configurations. Notice that we were not able to extract useful insights regarding the discrimination between DEG and NODEG patients from an analysis based only on the anatomical features, such as aortic orifice’s area, sinotubular junction’s diameter and ascending aorta’s curvature.

**Fig. 4:**
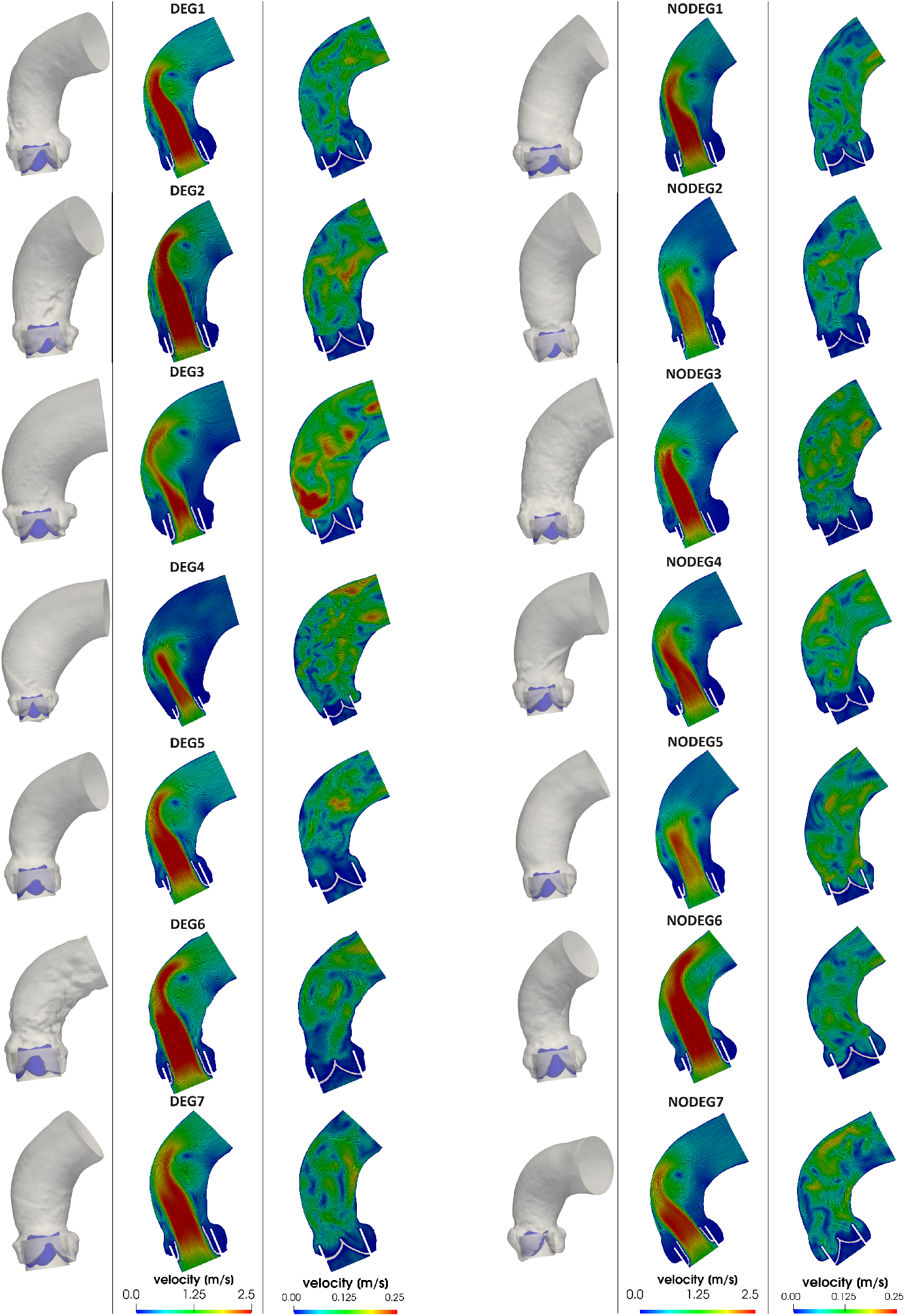
Computational domain with the bioprosthetic leaflets in open configuration (left); Velocity streamlines on a longitudinal section at the systolic-peak (middle) and a representative diastolic (right) instants.

In addition to the pre-operative CT scans available for each patient in the study population, for patient DEG1 and NODEG1 we also have at disposal post-operative CT scans taken few years after the intervention, which can be used to validate the virtual stent implantation procedure. In particular, starting from these post-operative CT images we reconstruct the post-operative aortic geometry and the implanted stent, and we compare the position of the latter with that obtained by the virtual insertion in the pre-operative scenario. Figure 5 shows the positions of the reconstructed (in red) and of the virtually inserted (in green) stents. We observe that the position and orientation of the virtual stent inside the pre-operative geometry is in qualitative good agreement with those of the implanted stent inside the post-operative geometry. Moreover, the reconstructed implanted stent (in red) shows a circular cross section in the view from the left ventricle, supporting our decision to consider a cylindrical virtual stent.

**Fig. 5:**
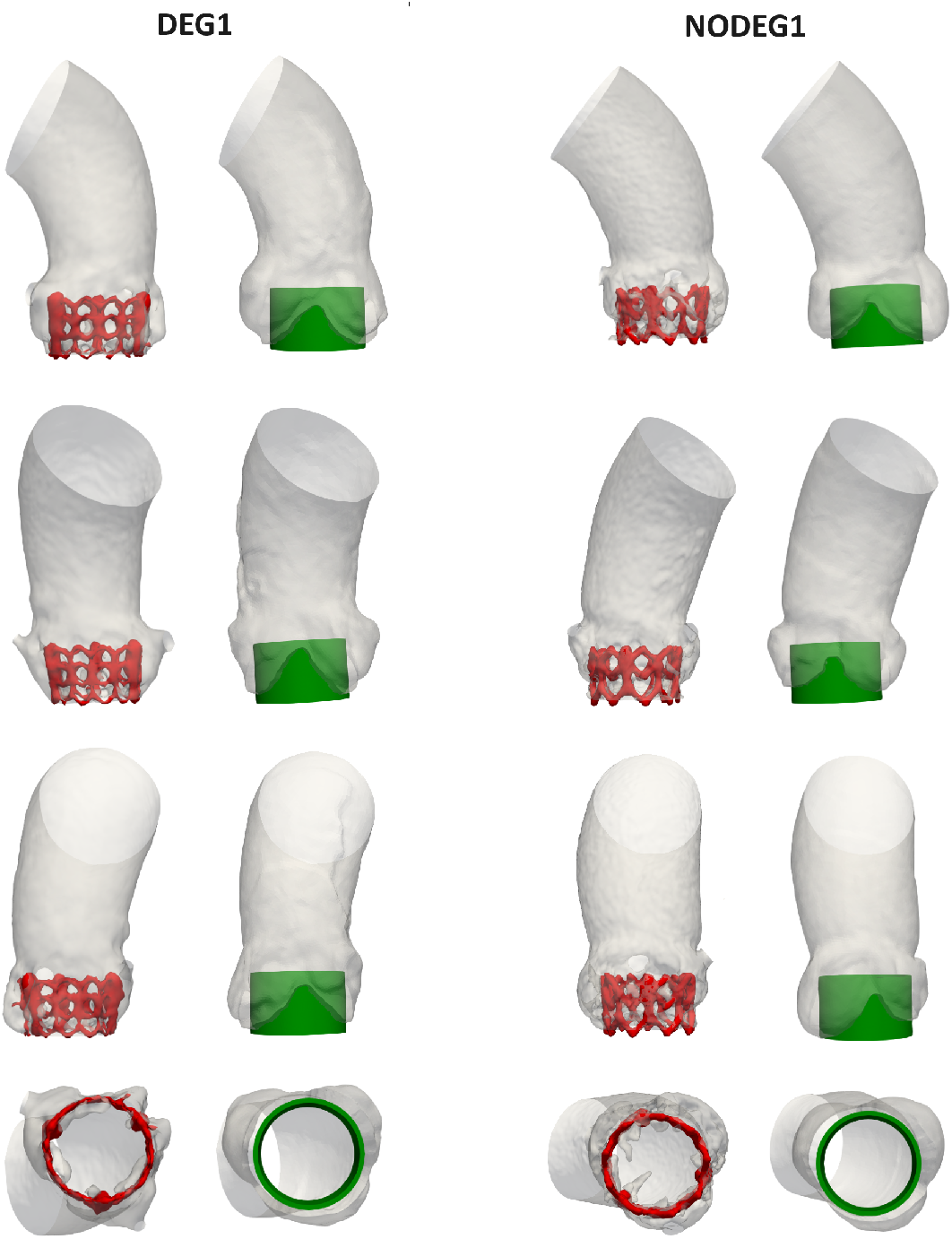
Validation of virtual stent implantation for patients DEG1 and NODEG1. Three different frontal views and the view from the left ventricle are reported in the different rows. Left: Aortic geometry and implanted stent (in red) reconstructed from the post-operative CT scan; Right: computational domain after virtual stent insertion (in green).

### 3.2 Analysis of the numerical results

In Figure 4, the blood velocity streamlines resulting from the CFD analysis are depicted on a longitudinal plane at two time instants: one during systole and one during diastole. The CFD results show that, during systole, the presence of the bioprosthetic TAVI valve results in the formation of a high velocity jet inside the ascending aorta together with some vortical structures. The morphology of the jet and of the vortical structures depends on the patient-specific geometries and cardiac outputs (see Table 2). The results at the diastolic time are characterized by chaotic velocity streamlines throughout the computational domain for each patient.

Together with this jet, also vortical structures collide against the aortic wall, giving rise to a highly disturbed flow and great shear stresses on the wall. This is reported for a representative case (patient DEG3) in Figure 6, where the WSS pattern is shown to follow the evolution of the vortical structures, which are highlighted by the Q-criterion. In the same figure we report also the pressure field at a time instant in early systole on a longitudinal plane. This result shows a relevant pressure drop across the bioprosthetic valve and low pressure zones in the ascending aortic tract, in correspondence of the vortices. Finally, we report also the hemodynamic indices presented in Section 2.5.

**Fig. 6:**
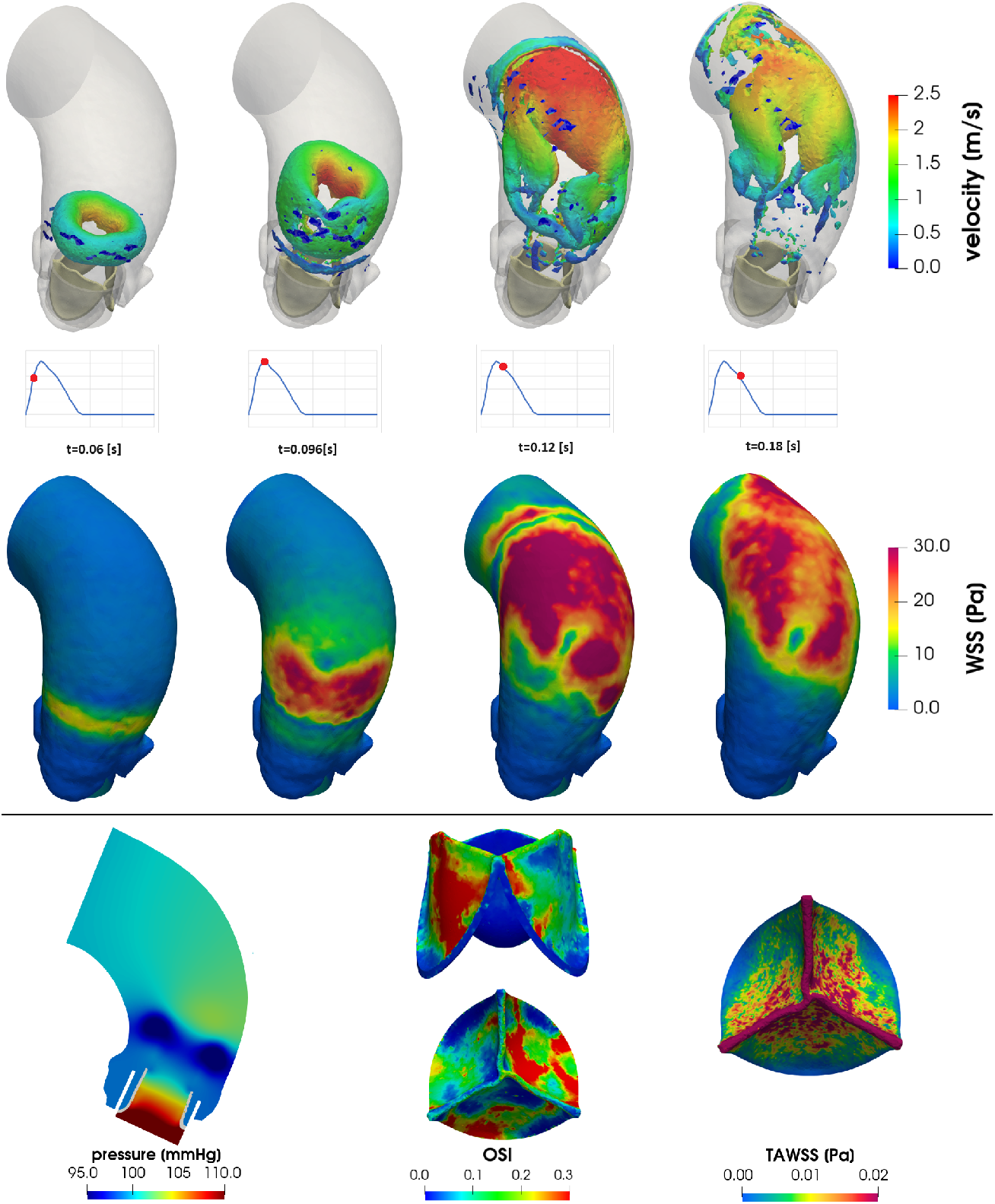
Post-processing of the numerical results for patient DEG3. Top row: time evolution of vortical structures, isolated exploiting the Q-criterion (*Q >* 50000 *Hz*^2^) and coloured by velocity. Middle row: time evolution of WSS on the aortic wall. Bottom row: pressure field at a time instant in early systole (left); OSI on the bioprosthetic leaflets during both systole and diastole (middle); diastolic TAWSS on the bioprosthetic leaflets (right).

### 3.3 Statistical Analysis

In Table 3 we report, for each patient, the hemodynamic scores introduced in Section 2.5. In Figure 7a we report the corresponding boxplots, which highlight that the scores are distributed differently in the DEG and NODEG patients. In particular, each hemodynamic index shows an increased variability (wider boxes) in the DEG patients with respect to the NODEG ones, with the only exception of *avgTAWSS* which takes low values for each DEG patient.

**Table 3:** Hemodynamic scores introduced in Section 2.5.

| ID | $\overline{A^{WSS}}$ | $\overline{V_Q^{\%}}$ | $avgOSI$ | $avgTAWSS$ |
| --- | --- | --- | --- | --- |
| **DEG1** | $1.48 \times 10^{-3}$ | $2.81 \times 10^{-2}$ | $1.43 \times 10^{-1}$ | $1.03 \times 10^{-2}$ |
| **DEG2** | $1.25 \times 10^{-3}$ | $3.68 \times 10^{-2}$ | $1.57 \times 10^{-1}$ | $1.21 \times 10^{-2}$ |
| **DEG3** | $1.48 \times 10^{-3}$ | $2.46 \times 10^{-2}$ | $1.61 \times 10^{-1}$ | $1.18 \times 10^{-2}$ |
| **DEG4** | $1.41 \times 10^{-3}$ | $1.30 \times 10^{-2}$ | $1.35 \times 10^{-1}$ | $0.98 \times 10^{-2}$ |
| **DEG5** | $1.16 \times 10^{-3}$ | $3.20 \times 10^{-2}$ | $1.65 \times 10^{-1}$ | $0.79 \times 10^{-2}$ |
| **DEG6** | $1.04 \times 10^{-3}$ | $4.18 \times 10^{-2}$ | $1.51 \times 10^{-1}$ | $0.91 \times 10^{-2}$ |
| **DEG7** | $0.96 \times 10^{-3}$ | $3.43 \times 10^{-2}$ | $1.94 \times 10^{-1}$ | $1.24 \times 10^{-2}$ |
| **NODEG1** | $1.20 \times 10^{-3}$ | $2.86 \times 10^{-2}$ | $1.43 \times 10^{-1}$ | $2.34 \times 10^{-2}$ |
| **NODEG2** | $0.92 \times 10^{-3}$ | $1.69 \times 10^{-2}$ | $1.31 \times 10^{-1}$ | $0.87 \times 10^{-2}$ |
| **NODEG3** | $1.30 \times 10^{-3}$ | $1.88 \times 10^{-2}$ | $1.50 \times 10^{-1}$ | $1.01 \times 10^{-2}$ |
| **NODEG4** | $1.24 \times 10^{-3}$ | $2.23 \times 10^{-2}$ | $1.33 \times 10^{-1}$ | $0.90 \times 10^{-2}$ |
| **NODEG5** | $1.00 \times 10^{-3}$ | $2.13 \times 10^{-2}$ | $1.30 \times 10^{-1}$ | $3.21 \times 10^{-2}$ |
| **NODEG6** | $1.29 \times 10^{-3}$ | $4.35 \times 10^{-2}$ | $1.29 \times 10^{-1}$ | $4.03 \times 10^{-2}$ |
| **NODEG7** | $1.27 \times 10^{-3}$ | $2.72 \times 10^{-2}$ | $1.41 \times 10^{-1}$ | $1.10 \times 10^{-2}$ |

**Fig. 7:**
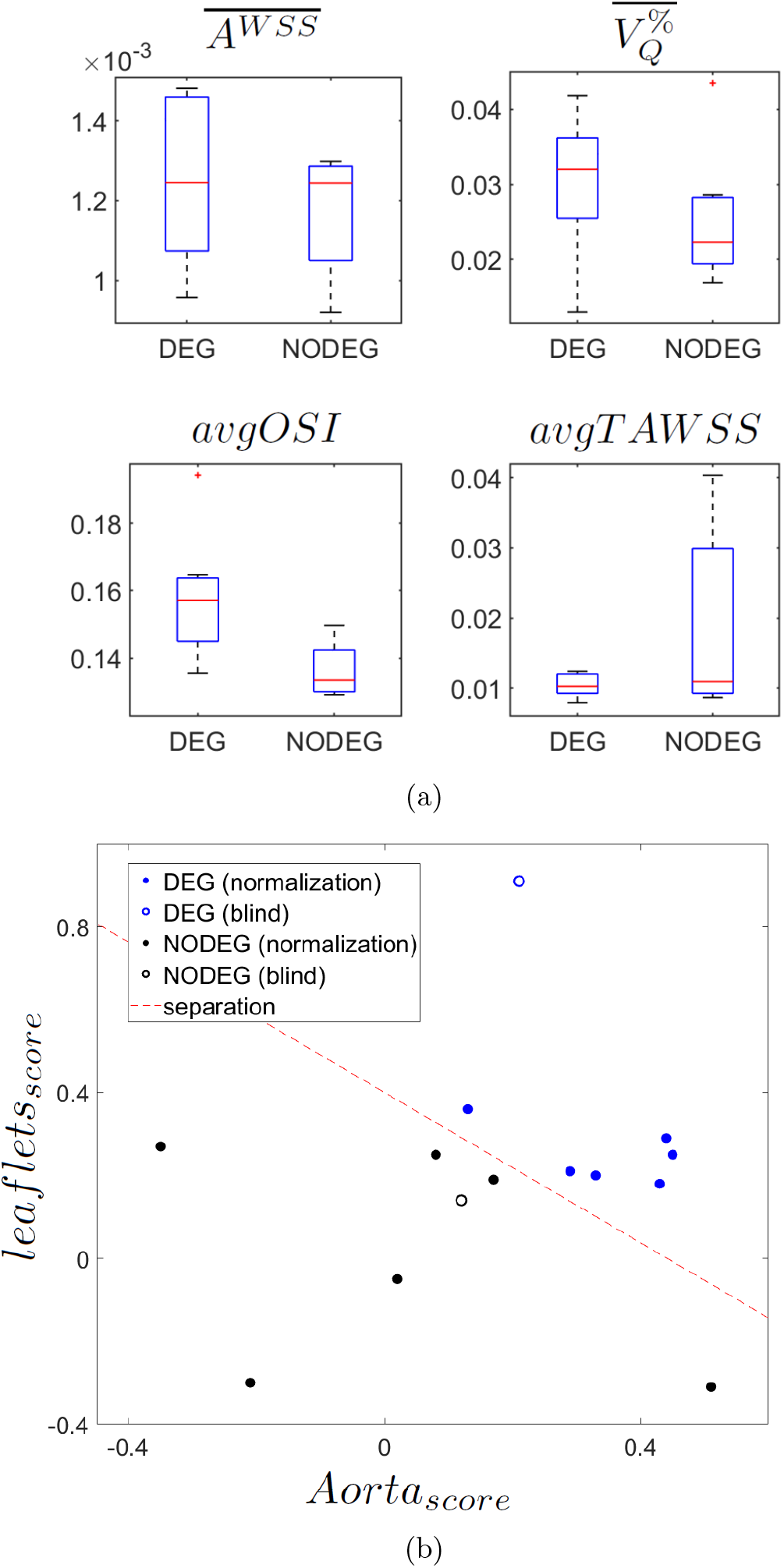
(a) Visualization of the hemodynamic scores distribution in the DEG and NODEG patients using boxplots. (b) Visualization of DEG and NODEG patients in a two-dimensional space given by *Aorta*_*score*_ and *leaflets*_*score*_.

To assess differences between the two subgroups of patients in terms of each individual hemodynamic score, the Wilcoxon rank-sum test is employed: we found that *avgOSI* is the only index showing statistically significant differences (*p* = 0.007) between DEG and NODEG patients, whereas for the other indices we have in any case *p >* 0.1. However, *avgOSI* alone is not able to clearly separate the two groups of patients. This is the reason why we focused on synthetic scores (see Section 3.4).

### 3.4 Synthetic scores

In Figure 7b we visualize each patient in a two-dimensional space using the synthetic scores *Aorta*_*score*_ and *leaflets*_*score*_ introduced in Section 2.5. The figure shows that DEG and NODEG patients can be linearly separated in this two-dimensional space. This result highlights that, by considering the synthetic scores instead of the individual ones, we are able to discriminate more efficiently the two groups of patients.

We report in Table 4, for both the normalization and blind sets, the global *SV D*_*score*_ introduced in Section 2.5. In particular, for the normalization set DEG patients present *SV D*_*score*_ values greater than or equal to 0.15, whereas NODEG patients present a score lower than 0.15. Thus, this global synthetic score seems to be able to discriminate between the two subgroups of patients and can be interpreted as likelihood of developing a premature onset of SVD. This is confirmed by the application of this score to two cases (DEG7 and NODEG7) not used during the normalization. Specifically, as reported in Table 4 we obtained *SV D*_*score*_ = 0.56 for DEG7 and *SV D*_*score*_ = 0.09 for NODEG7.

**Table 4:** Global SVD-discriminating score *SVD*_*score*_. Patient DEG7 and NODEG7 belong to the blind set.

| ID | $SVD_{score}$ | ID | $SVD_{score}$ |
| --- | --- | --- | --- |
| **DEG1** | 0.29 | **NODEG1** | -0.06 |
| **DEG2** | 0.23 | **NODEG2** | 0.04 |
| **DEG3** | 0.31 | **NODEG3** | 0.11 |
| **DEG4** | 0.15 | **NODEG4** | 0.12 |
| **DEG5** | 0.27 | **NODEG5** | -0.34 |
| **DEG6** | 0.30 | **NODEG6** | 0.11 |
| **DEG7** | 0.56 | **NODEG7** | 0.09 |

## 4 Discussion

Structural Valve Deterioration (SVD) is a complex phenomenon whose underlying mechanisms are still incompletely understood. Recent studies suggest the host’s immune response as a major factor of SVD pathogenesis, manifested by a combination of processes phenocopying atherosclerosis and calcification of native aortic valves [10]. Moreover, calcific aortic valve disease is an active process characterized by lipoprotein deposition and chronic inflammation [47]. For these reasons we believe that aortic hemodynamics can have an impact on the development of SVD. In particular, we investigate blood-dynamics features also downstream the valve because we are looking into possible correlations with SVD, even if not directly causing it.

Notice that in this study we exploit only data routinely acquired during the diagnosis and treatment of Aortic Stenosis (AS), such as pre-operative CT scans and post-operative TTE.

In conclusion, we can state that the identification of a hemodynamic score (*avgOSI*) that shows statistically significant differences (*p* = 0.007) between DEG and NODEG patients and the definition of purely hemodynamic-based synthetic scores able to discriminate between the two subgroups of patients suggest that post-operative hemodynamics should be taken into account for a complete assessment of TAVI bio-prosthetic valves long-term durability. As a consequence, the proposed hemodynamic indices, possibly together with other SVD predictors [3], can assist clinicians in a patient-specific planning of follow-up exams based on the risk of prematurely developing SVD. Specifically, patients who are predicted to encounter a premature onset of SVD could be monitored in a systematic way by the hospital where TAVI is performed in order to avoid the dispersion of the patients, which represents a strong limit of the follow-up procedure.

The main modeling limitations of this work are:

### Neglecting the wire-frame design of the stent

This is accurate only on the stent’s ventricular side, due to the presence of the inner skirt. This limitation could potentially have an impact on secondary diastolic flows which may cross the wire-frame and thus, in particular, on the leaflets’ OSI and TAWSS scores. We plan to provide a wire-frame design for the aortic side of the stent in future studies;

### Including patient-specific information concerning only mean flow rates in the boundary conditions

Prescribing completely patient-specific inlet conditions can be achieved by exploiting 4D-Flow Magnetic Resonance Imaging [48]. This could not be done in the present work due to the retrospective nature of the study. Alternatively, one could build more realistic 4D velocity profiles using, e.g., the techniques presented in [49].

### Not considering the interaction between fluid and the prosthesis

This is done according to the results of the preliminary study [11], where no significant indices were identified from this interaction;

### Not accounting for coronary flow

Coronary flow is thought to mainly come from the elastic relaxation of the aortic wall following its systolic expansion [50, 51], but we cannot describe this process due to the rigid wall assumption in the proposed CFD setting (see Section 2.3);

### Analyzing only fourteen patients

We understand that this number is probably too small to provide significant answers in terms of the influence of hemodynamics on SVD; however, collecting a greater number of data related to degenerated cases with available follow-up is difficult due to both the relative young age of the TAVI procedure and the high dispersion of patients after the implant;

### Using the same heart rate for all the patients due to the absence of such data for our cases

This may have an influence on fluid dynamics patterns and thus on the proposed hemodynamic scores. This deserves further and deeper investigations.

We remember that our model is subject to three main modeling assumptions which were made in order to reduce the computational effort and to make our analysis potentially reliable for clinical purposes:

- Representing the stent with an *a priori* -decided cylindrical shape without performing an inflation mechanical simulation (the latter was performed for example in [52]). We believe that our assumption is reasonable since, after the deployment, balloon-expandable valves are expected to achieve a cylindrical shape [53];
- Neglecting leaflets’ dynamics and leaflets’ Fluid-Structure Interaction. We model the valve’s leaflets in an on/off modality by assuming rigid systolic and diastolic configurations. This prevents us from accounting for the opening and closure mechanisms which may have an impact on the systolic jet and diastolic vortices. Moreover, we could not account for the fluttering of the open leaflets during systole. However, we believe that using a realistic effective opening area for the bioprosthetic valve, as done in this work, is the most influential feature, related to valve modeling, to accurately describe post-TAVI aortic hemodynamics, since this allows us to achieve realistic velocity and stress magnitudes inside the aorta;
- Describing the aortic wall as rigid. We accept this simplification since we are analyzing elderly patients often showing calcification in the ascending aorta, thus we expected a limited elasticity of the vessel wall.

## Declarations

### Ethics approval

The study was approved from the Institutional Review Board of Centro Cardiologico Monzino and registered with number R1264/20. Informed consent was obtained from all patients.

## Acknowledgments

LC, CV, IF are members of the INdAM group GNCS “Gruppo Nazionale per il Calcolo Scientifico” (National Group for Scientific Computing). CV has been partially supported by the Italian Ministry of University and Research (MIUR) within the PRIN (Research projects of relevant national interest) MIUR PRIN22-PNRRn. P20223KSS2 “Machine learning for fluid-structure interaction in cardiovascular problems: efficient solutions, model reduction, inverse problems, and by the Italian Ministry of Health within the PNC PROGETTO HUB - DIAGNOSTICA AVANZATA (HLS-DA) “INNOVA”, PNC-E3-2022-23683266.

